# Mini-agrin prevents calcium leakage and restores the dystrophin complex

**DOI:** 10.1101/2025.11.25.690165

**Authors:** Hitham Aldharee, Bartosz Mierzejewski, Alexander Kerr, Laricia Bragg, Anne Bigot, Judith R. Reinhard, Sabrina Santoleri, Neil Roberts, Raman Das, Vincent Mouly, Marcus A. Ruegg, Giulio Cossu, Francesco Galli

## Abstract

Muscle cell death in muscular dystrophies depends upon calcium ion (Ca^++^) leakage through sarcolemma and sarcoplasmic reticulum, triggered by muscle stretch during eccentric contraction.

We show here that Ca^++^ spikes are detected in dystrophic myogenic cells in culture since early differentiation, before sarcomere assembly and contraction. Healthy and genetically corrected dystrophic myotubes do not display Ca^++^ spikes which are blocked by co-culturing DMD myogenic cells with embryonic mouse motoneurons or treating them with agrin proteoglycan. Same effect is elicited by a muscle spliced, COOH peptide of agrin (termed here mini-agrin) that interacts with dystroglycan, favouring its binding to the basal lamina.

Lack of dystrophin in DMD myotubes results in decreased expression of Ca_V_1.1 (CACNA1S), a Ca^++^ sensor component of the Dihydropyridine Receptor (DHPR) complex, known to regulate Ryanodine Receptor 1 (RyR1). These events explain the emergence of Ca^++^ spikes. Mini-agrin addition to medium, or lentivector-mediated mini-agrin expression in transplanted cells in vivo, stabilize the expression of Ca_V_1.1 on the membrane. This leads to disappearance of Ca^++^ spikes and to reappearance of α-dystroglycan, α-sarcoglycan and n-NOS, indicating the reconstitution of the dystrophin complex in the absence of dystrophin. These findings unveil a novel regulatory mechanism and offer a new therapeutic opportunity for targeting calcium ion influx as a co-treatment strategy.

## Introduction

Muscular dystrophies (MD) are genetic disorders that affect skeletal muscle and often the heart^1,2^. These conditions vary in age of onset, severity, and muscles affected. MD still lack effective treatments. Currently, steroids are the standard therapy, but they only slow disease progression and cause significant side effects. Many MDs, including Duchenne Muscular Dystrophy (DMD; OMIM #310200), result from mutations in genes that encode structural proteins forming the dystrophin-glycoprotein complex (DGC), which connects the muscle cell cytoskeleton to the basal lamina^2,3^. Dystrophin, the protein produced by the DMD gene, plays a central role in this complex. Mutations disrupting the reading frame cause DMD, while in-frame deletions lead to the milder Becker Muscular Dystrophy (BMD)^4,5,6^. The process by which the absence of dystrophin results in micro-tearing of the membrane and/or irregularities in ion channels, particularly Ca^++^ channels, is not yet fully understood. However, it is well known that Ca^++^ leaks into the myofibre cytosol from both the sarcolemma and the sarcoplasmic reticulum, initiating a cascade involving activation of Ca^++^-dependent proteases that ultimately cause muscle fibre death. Few clinical trials using different Ca^++^ inhibitors gave inconclusive results, likely because of limited patient numbers and systemic side effects^7^. In this study, we examined the effect of dystrophin absence on differentiating muscle cells in culture. We used immortalised human wild-type (WT) and DMD myogenic cells carrying a skippable mutation in exon 51 of the dystrophin gene^8^, some of which were genetically corrected using a lentivector expressing U7 snRNA (DMD-U7)^9^. We also extended our investigation to primary DMD mesoangioblasts (Mabs) and their genetically corrected counterparts.

We observed that both immortalized DMD myogenic cells and primary DMD Mabs exhibit transient Ca^++^ spikes, detected using Fura-2 labelling, which are absent in WT cells. Importantly, genetic correction leading to dystrophin re-expression abolishes these Ca^++^ spikes. Moreover, co-culture with mouse embryonic motoneurons or the administration of agrin alone also eliminates Ca^++^ spikes. We show here that agrin stabilises the Ca_V_.1.1 sensor channel, which regulates the Ryanodine Receptor complex and ultimately prevents Ca^++^ spikes. These results unveil a novel and early mechanism of Ca^++^ dysregulation in DMD myogenic cells and suggest a novel potential therapeutic tool. Specifically, a COOH agrin peptide^10^, here referred to as mini-agrin, is sufficient to block Ca^++^ influx in DMD cells and reconstitute the DGC in the absence of dystrophin. This approach holds promise for addressing the dysregulated Ca^++^ signalling implicated in myofibers degeneration and may offer a novel therapeutic strategy for DMD and other MDs.

## Results

### Lack of dystrophin causes Ca^++^ leakage in early differentiated DMD myotubes

We observed that non-innervated, early differentiated DMD myotubes display random, repetitive spikes, detected by the Ca^++^ indicator Fura-2, whereas WT myotubes do not (Fig.1A and Suppl. Movies 1,2). Interestingly, DMD myotubes, genetically corrected with the U7snRNA (DMD-U7)^1^, also show no detectable Ca^++^ spikes (Fig.1A and Suppl. Movie 3), indicating that the expression of dystrophin is sufficient to prevent the occurrence of Ca^++^ spikes. These findings align with previous observations in *mdx* mice and human DMD myofibres^11,12^.

**Fig. 1:**
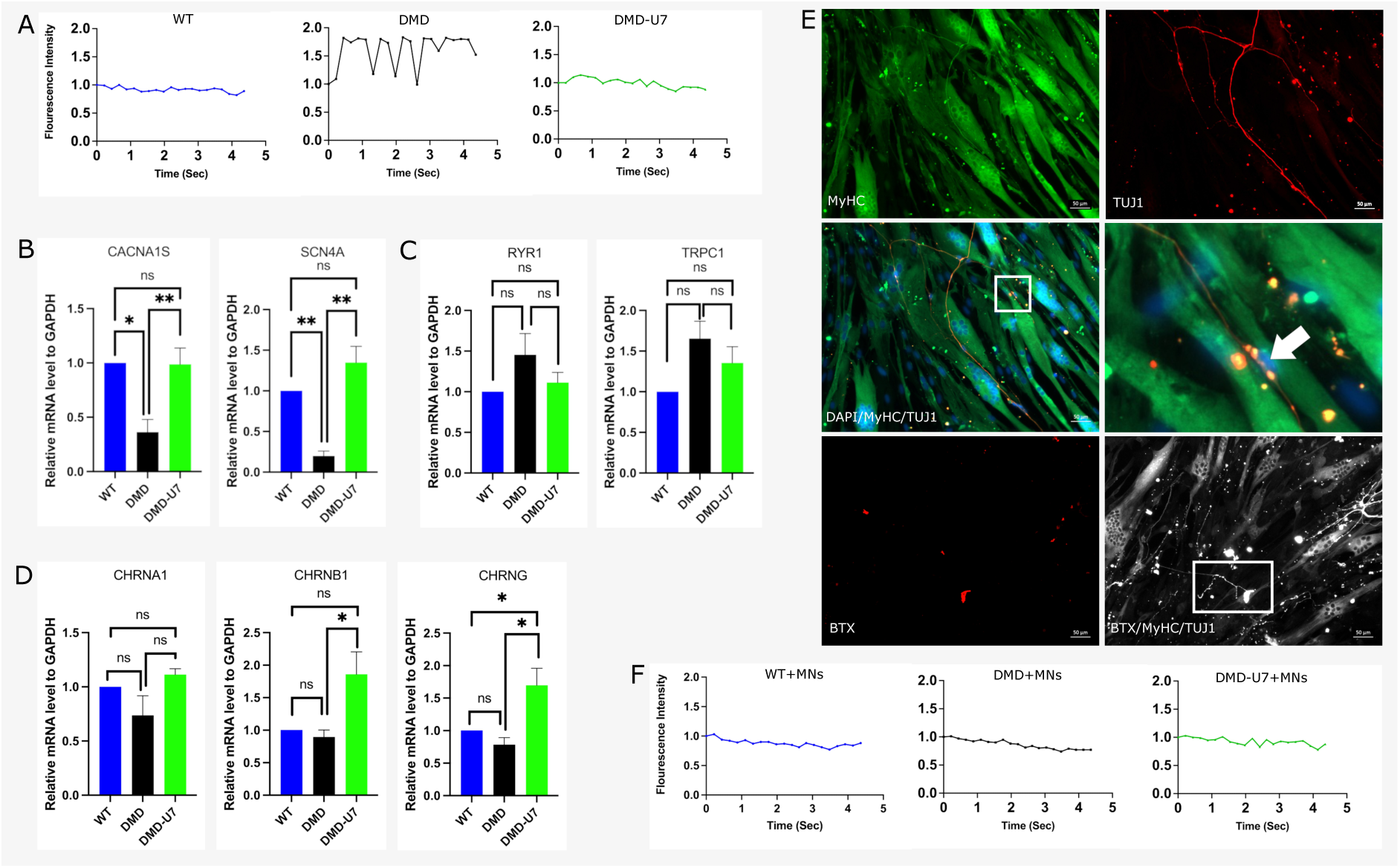
Early Dystrophin Deficiency Impairs Calcium Handling and Muscle Contraction in DMD Myotubes. **A.** Representative fluorescence intensity (FI) analysis over time in WT, DMD, and DMD U7 myotubes. Values were normalized to FI at time 0 (n=3). **B.** qRT-PCR analysis of relative mRNA expression of **CACNA1S** and **SCN4A** in WT, DMD, and DMD-U7 myotubes (n=3, *p<0.05). **C.** qRT-PCR analysis of relative mRNA expression of **RYR1** and **TRPC1** in WT, DMD, and DMD U7 myotubes (n=3, *p<0.05). **D.** qRT-PCR analysis of relative mRNA expression of α1 (CHRNA1), β1 (CHRNB1) and γ (CHRNG) acetylcholine subunits in WT, DMD, and DMD-U7 myotubes (n=3, *p < 0.05). **E.** Immunofluorescence staining of DMD myotubes co-cultured with motoneurons (MNs). Myotubes were labeled with α-Myosin Heavy Chain (MyHC, green), MNs with α-TUJ1 (red), and acetylcholine receptors (AChR) with α-bungarotoxin (BTX, red) to show neuromuscular junction formation (n=3). **F.** Representative FI analysis over time in WT, DMD, and DMD-U7 myotubes co-cultured with MNs. Values were normalized to FI at time 0 (n=3).

Additionaly, myotubes derived from DMD mesoangioblasts (Mabs) also show transient Ca^++^ spikes that are absent in both myotubes derived from WT Mabs and in myotubes derived from genetically corrected DMD Mabs (Suppl. Fig.1A), further reinforcing the role played by dystrophin in preventing Ca^++^ spikes.

### Calcium regulatory genes are downregulated in DMD myotubes

To investigate whether the alteration of Ca^++^ homeostasis in DMD myotubes could be attributed to changes in the expression of genes regulating ion channels or sensors, we performed qRT-PCR analysis to assess their relative mRNA levels. We observed a dramatic reduction in the expression of both calcium voltage-gated channel subunit α1S (CACNA1S) gene and voltage-gated channel α4 subunit (SCN4A) gene in DMD myotubes (Fig.1B); the first encodes the skeletal muscle L-type voltage-gated Calcium Channel α1 subunit of the DHPR (Ca_V_1.1) and the second a sodium channel. Of note, these genes were expressed in DMD-U7 myotubes at levels comparable with WT myotubes (Fig.1B). It is known that the DGC constitutes a scaffold for various channel proteins involved in signalling processes ^13^. This suggests that the absence of the dystrophin could be responsible for the observed altered expression of both SCN4A and CACNA1S. Conversely, RyR1 and transient receptor potential cation channel subfamily C (TRPC1) showed slight upregulation in DMD myotubes, with no significant changes in DMD-U7 myotubes compared to healthy myotubes (Fig.1C). Additionally, we found that both α1 (CHRNA1), β1 (CHRNB1) and γ (CHRNG) subunits of the AChR were modestly reduced in DMD myotubes (Fig. 1D).

The L-type Voltage Gate Calcium Channel (VGCC), Ca_V_1.1 isoform, is localized on sarcolemma and T-tubules^14^, where it interacts with and modulates the RyR1^15,16^. Ca_V_1.1 is only expressed and tightly regulated in skeletal muscle^17,18^. Previous studies indicated that Ca_V_1.1 function is altered in muscle diseases such as DMD^11,18,19^. To determine whether Ca_V_1.1 contributes to the observed Ca^++^ leakage, DMD myotubes were treated with specific calcium channel blockers, nimodipine and nitrendipine, which effectively abolished the Ca^++^ spikes (Suppl. Fig.1B). Caffeine-dependant-Ca^++^ release from RyR1 was larger in DMD than in WT myotubes, even if the difference was not statistically significant (Suppl. Fig.1B).

### Innervation suppresses Ca^++^ spikes in DMD myotubes

Under our culture conditions, Ca^++^ spikes were not followed by muscle contraction, most likely due to the absence of a developed and mature contractile apparatus. To confirm this hypothesis, we analysed the level of maturation in WT, DMD and DMD-U7 myotubes using transmission electron microscopy (TEM) (Suppl. Fig.2A). We observed initial assembly of thick and thin filaments and early forming Z lines in all the different myotubes but no mature sarcomeres which explains why we did not observe any twitching myotube despite Ca^++^ spikes. To promote maturation, we co-cultured myotubes with freshly isolated motoneurons from E14 mouse embryonic spinal cord, embedded in a layer of Matrigel that separated them from underlying myotubes. Bungarotoxin staining of the co-culture indicated the formation of neuro-muscular junctions (NMJ) (Fig.1E), confirmed by the TEM finding of advanced sarcomerogenesis (Suppl Fig.2B), even though myotubes were still unable to contract. Unexpectedly, we observed the disappearance of the Ca^++^ spikes in innervated DMD myotubes, even if motoneurons were not stimulated. No Ca^++^ spikes were observed, in both healthy and DMD-U7 myotubes under these conditions (Fig.1F) (Suppl Movies 4,5,6).

**Fig. 2:**
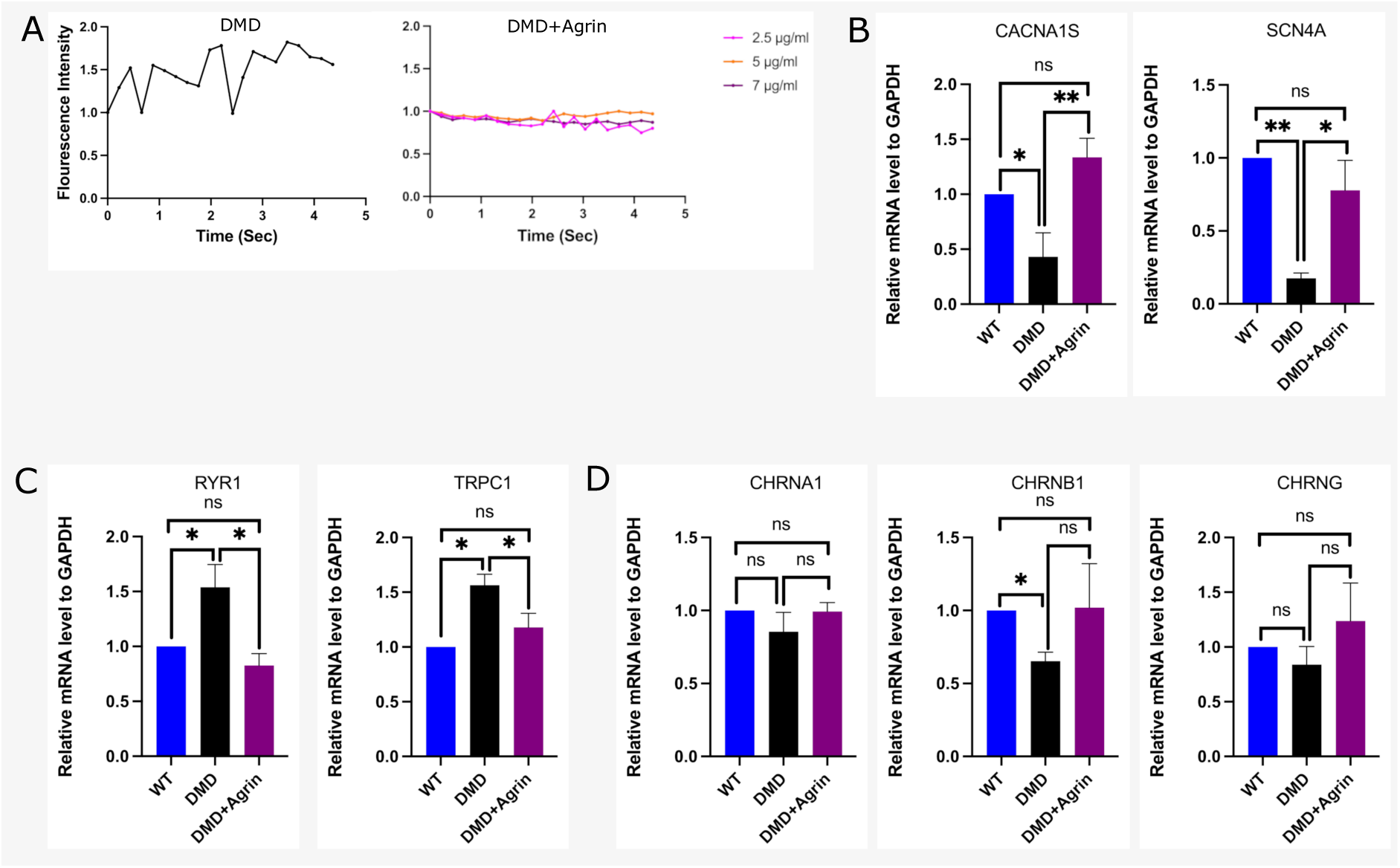
Agrin Restores Calcium Homeostasis and Gene Expression in DMD Myotubes. **A.** Representative fluorescence intensity (FI) analysis over time in DMD, and DMD myotubes treated with different concentration of Agrin (2.5 μg/ml, 5 μg/ml, 7 μg/ml). Values were normalized to FI at time 0 (n=3). **B.** qRT-PCR analysis of relative mRNA expression of **CACNA1S** and **SCN4A** in WT, DMD, and DMD treated with Agrin (7 μg/ml) myotubes (n=3, *p<0.05). **C.** qRT-PCR analysis of relative mRNA expression of **RYR1** and **TRPC1** in WT, DMD, and DMD treated with Agrin (7 μg/ml) (n=3, *p<0.05). **D.** qRT-PCR analysis of relative mRNA expression of α1 (CHRNA1), β1 (CHRNB1) and γ (CHRNG) acetylcholine subunits in WT, DMD, and DMD treated with Agrin (7 μg/ml) myotubes (n=3, *p < 0.05).

The disappearance of Ca^++^ spikes in the absence of motoneuron stimulation, suggested that molecules released at the NMJ level may be responsible for this effect. Given its role in the development of the NMJ and in promoting binding to the basal lamina, agrin, a heparate sulfate proteoglycan relased by both nerve and muscle, was an obvious candidate molecule to be tested.

### Agrin abolishes Ca^++^ spikes and restores normal expression of calcium regulatory genes in DMD myotubes

To investigate whether agrin directly influences calcium signaling in DMD muscle, we analysed its effect on spontaneous Ca^++^ activity and the expression of key genes in Ca^++^ homeostasis in DMD myotube. When added to cultures of DMD myotubes without innervation, agrin, at any concentration tested in the μM range, abolished Ca^++^ spikes in DMD myotubes, indicating that the effect of muscle innervation can be explained by agrin release (Fig.2A, Suppl. Movies 7,8,9). To understand how agrin abolishes the spikes, we investigated whether it affects the expression of genes involved in Ca^++^ homeostasis. We observed that gene expression of CACNA1S and SCN4A, decreased in DMD myotubes, restored to normal levels in agrin-treated DMD myotubes (Fig.2B). We also found that both RyR1 and TRPC1 were expressed at similar level in all samples (Fig.2C) while both α1 (CHRNA1), β1 (CHRNB1) and γ (CHRNG) subunits of the AChR were not significantly modified by agrin (Fig.2D). These results show that agrin may interfere with Ca^++^ homeostasis by modifying related genes expression, in skeletal muscle, such as CACNA1S and SCN4A.

### Agrin is essential for Ca_V_1.1 expression

Expression of Ca_V_1.1 protein (CACNA1S) is downregulated in *mdx* mice^11,18^. Here, western blot analysis showed that Ca_V_1.1 expression was significantly reduced in DMD myotubes but restored in DMD-U7 myotubes (Fig.3A). This result confirms observations at the mRNA level, suggesting that the absence of dystrophin negatively affects Ca_V_1.1 expression at very early stages of differentiation. Remarkably, Ca_V_1.1 expression was successfully restored in DMD myotubes treated with agrin, despite the continued absence of dystrophin (Fig.3B). Interestingly, WB analysis revealed that agrin treatment in DMD myotubes also restored dystroglycan expression (Suppl. Fig.3).

**Fig. 3:**
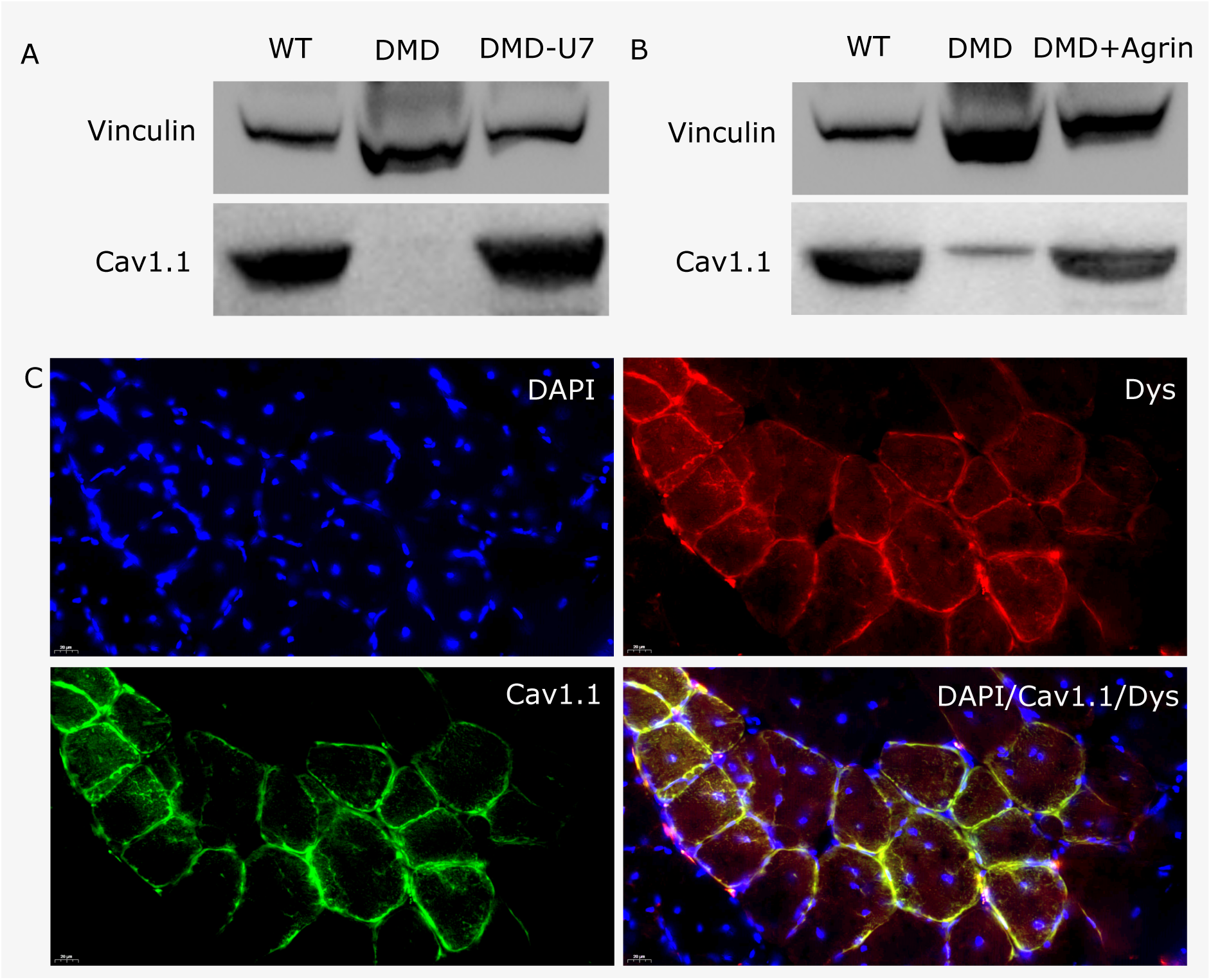
Agrin Restores Ca_V_1.1 Expression *in vitro* and *in vivo*. **A.** Representative Western blot (WB) showing Ca_V_1.1 expression in WT, DMD, and DMD-U7 myotubes (n = 3). **B.** Representative WB showing Ca_V_1.1 expression in WT, DMD, and DMD myotubes treated with agrin (7 μg/ml) (n=3). **C.** Immunofluorescence (IF) images of transverse sections of tibialis anterior (TA) muscles from NSG-mdx-Δ51 mice transplanted with 5 × 10⁵ WT cells, showing expression of dystrophin (red) and Ca_V_1.1 (green) (n=3).

To confirm the role of dystrophin in regulating Ca_V_1.1 expression *in vivo*, we analysed by IF the *Tibialis Anterior* of NSG-DMD mice previously transplanted with DMD-U7 myoblasts. We found that only dystrophin expressing myofibers show robust expression of Ca_V_1.1 which is barely detectable in DMD non corrected myofibers (Fig.3C).

### Mini-Agrin restores Ca_v_1.1 expression both in vivo and in vitro

Agrin is a large molecule and cannot be not be veicolated through a viral vector. However, it was shown in the past that the COOH terminal region of agrin contains two differentially spliced peptides, one present in the version secreted by motoneurons and involved in clustering the AchR, the other (C95 0,0) present in the version secreted by muscle and involved in binding to dystroglycan and laminin^20^. To test the role of this peptide (termed “mini-agrin”) in regulating the expression of Ca_V_1.1, we generated a lentivector encoding mini-agrin. The mini-agrin cDNA includes, after a signal peptide, the domain (C95 0,0) required for binding to α-dystroglycan but lacks the domain to bind α7β1 integrin (Suppl. Fig.4A). Mini-agrin was previously shown to ameliorate muscle integrity in laminin-α2-deficient mice^10^. We then transduced DMD myogenic cells to constitutively express mini-agrin (DMD-mAgr) to test whether mini-agrin expression could modulate Ca_V_1.1 levels and suppress Ca^++^ transients. As a first step, we confirmed by immunofluorescence analysis that Ca_V_1.1 expression was restored in DMD-mAgr myotubes despite the absence of dystrophin (Suppl. Fig.4B). Using the Fura-2 assay, we observed that mini-agrin expression abolished Ca^++^ spikes in DMD-mAgr myotubes (Fig.4A) (Suppl. Movies 10). Subsequent immunofluorescence analysis showed that SERCA (sarcoplasmic/endoplasmic reticulum calcium ATPase), a key regulator of intracellular calcium homeostasis in muscle, was not altered in DMD-mAgr cells. Moreover, SERCA levels in DMD-mAgr myotubes were comparable to those in DMD cells (Suppl. Fig.4C). To confirm the ability of mini-Agrin to restore Ca_V_1.1 expression in vivo, we transplanted DMD-mAgr myogenic cells in the *Tibialis Anterior* of NSG-DMD. One month after transplantation, immunofluorescence analysis revealed that Ca_V_1.1 expression was also restored in vivo in the myofibres derived from DMD-mAgr myogenic cells (Fig.4B). Unexpectedly, myofibres expressing mini-Agrin also restored expression of α-sarcoglycan (Fig.4C) and nNOS (Suppl Fig.4D), indicating partial reassembly of the dystrophin associated complex (DGC) and recovery of muscle membrane function. We also observed by immunofluorescence analysis that mini-Agrin expression was not restricted to the areas where DMD-mAgr myogenic cells had engrafted (Lam AC–positive cells), but was also detected in regions lacking transplanted human cells (Fig.4D). Likewise, dystroglycan expression was not confined to areas containing DMD-mAgr myogenic cells but was also observed in resident mouse myofibres (Fig.4E). These findings suggest at least partial restoration of the DGC complex by mini-Agrin and indicate its ability to diffuse throughout skeletal muscle (likely between the sarcolemma and the basal lamina) and associate with the membrane of resident myofibers. These findings were further supported by WB analysis, which confirmed the expression of Ca_V_1.1, α-sargoclycan and α-dystroglycan in the *Tibialis Anterior* transplanted with DMD-mAgr myoblasts (Fig.4F, 4G). Altogether, these results highlight the role of agrin in regulating Ca^++^ homeostasis genes in skeletal muscle independently of dystrophin expression.

**Fig. 4:**
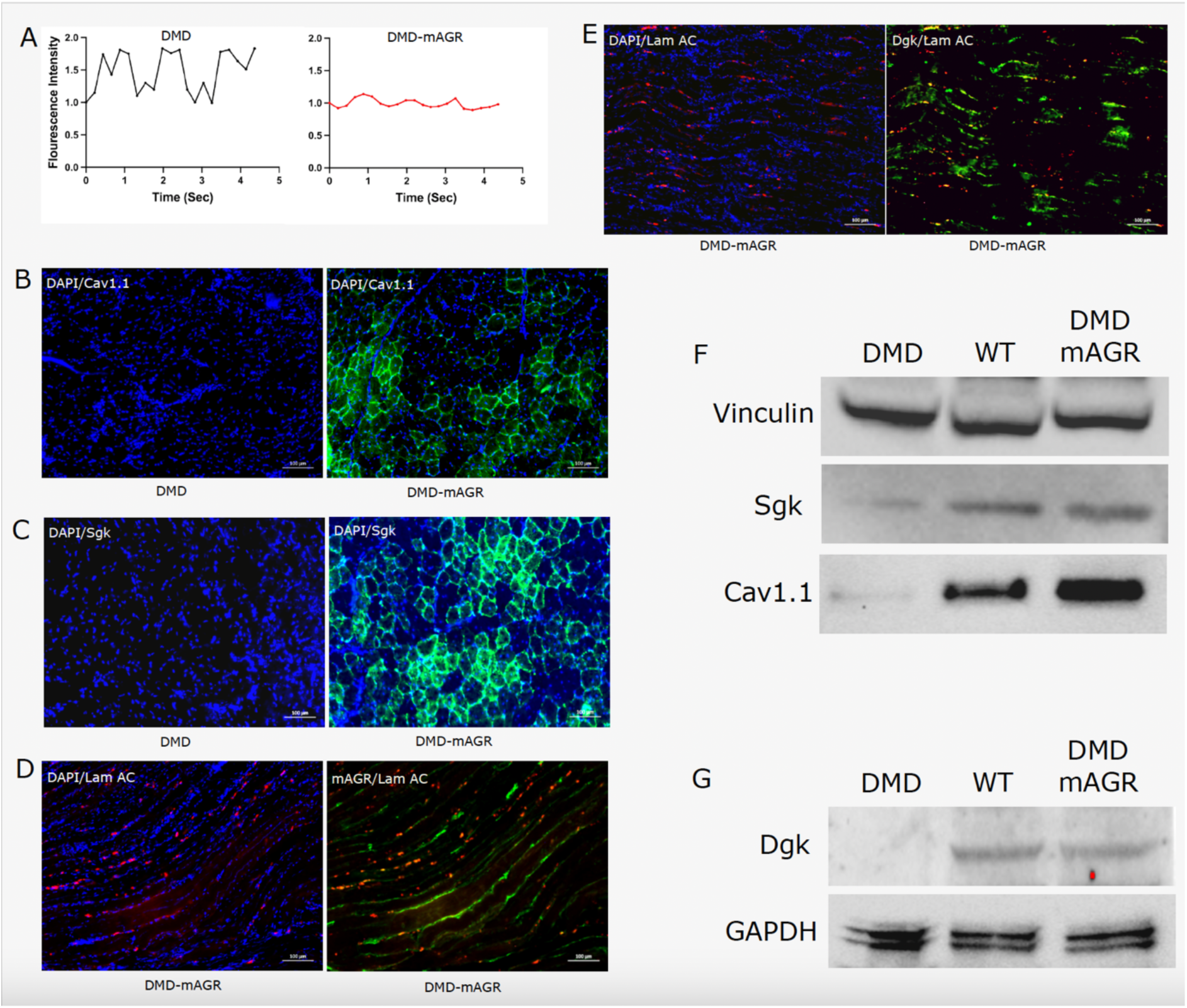
Mini-agrin Restores Ca_V_1.1 Expression in DMD Myotubes and in Vivo. **A.** Representative fluorescence intensity (FI) analysis over time in DMD and DMD-mAGR myotubes. Values were normalized to FI at time 0 (n=3). **B.** IF images of transverse sections of tibialis anterior (TA) muscles from NSG-mdx-Δ51 mice transplanted with 5 × 10⁵ DMD-mAGR cells, showing CaV1.1 expression (green) (n=3). **C.** IF images of transverse sections of TA muscles from NSG-mdx-Δ51 mice transplanted with 5 × 10⁵ DMD-mAGR cells, showing sarcoglycan (Sgk) expression (green) (n=3). **D.** IF images of longitudinal section of tibialis anterior (TA) muscles from NSG-mdx-Δ51 mice transplanted with 5 × 10⁵ DMD-mAGR cells,, showing Lamin AC (Lam AC) expression (red) and mini-Agrin (mAgr) expression (green) (n=3). **E.** IF images of longitudinal section of tibialis anterior (TA) muscles from NSG-mdx-Δ51 mice transplanted with 5 × 10⁵ DMD-mAGR cells,, showing Lam AC expression (red) and dystroglycan (Dgk) expression (green) (n=3). **F.** Representative Western blot (WB) showing CaV1.1 and sarcoglycan (Sgk) expression in total protein extracts from TA muscles of NSG-mdx-Δ51 mice either non-transplanted or transplanted with 5 × 10⁵ WT or DMD-mAGR cells (n=3). **G.** Representative WB showing dystroglycan (Dgk) expression in total protein extracts from TA muscles of NSG-mdx-Δ51 mice either non-transplanted or transplanted with 5 × 10⁵ WT or DMD-mAGR cells (n=3).

## Discussion

Although the concept that Ca^++^ leakage into DMD muscle fibres leads to degeneration has been established for decades, the exact mechanism remains incompletely understood. Indeed, Ca^++^ inhibitors have been tested as potential therapeutical treatments in patients, but results were inconclusive, partly due to low numbers of patients enrolled and the lack of specificity of these drugs for striated muscle^7^.

In this study, we observed frequent and repeated Ca^++^ spikes in two types of human DMD myogenic cells, but not in their healthy counterparts. Although this phenomenon had been previously observed^12^, the underlying mechanism had not been examined. Since spikes occur before sarcomerogenesis is completed, we could study the phenomenon independently of any mechanical effects caused by contraction. When cell-mediated exon skipping^9^ restored dystrophin expression, Ca^++^ spikes disappeared, in agreement with a previous report showing that expressing mini dystrophin in *mdx* mice resulted in a similar outcome^21^.

Co-culture with embryonic mouse motoneurons caused Ca^++^ spikes disappearance without activation of contraction, suggesting that secretory rather than electrical activity was responsible for the effect. Agrin appeared as an obvious candidate, given its role in regulating acetylcholine receptor assembly and other functions such as binding α-dystroglycan^22^ and establishing a connection with the basal lamina, as observed in congenital muscular dystrophy MDC1A caused by a mutation in laminin 211. Although agrin is a large heparan sulphate proteoglycan, making its therapeutic application challenging, the mini-agrin used here is a much smaller molecule that can be easily incorporated into a viral vector. Mini-agrin is currently being explored as a therapeutic approach for MDC1^10^.

We also observed that the expression of the Ca^++^ sensor Ca_V_1.1 was significantly reduced in DMD myotubes but restored in genetically corrected cells. Ca_V_1.1 is the only Ca^++^ sensor isoform expressed in skeletal muscle^15,16^ and it is an integral component of the DHPR complex, which regulates fluxes through RyR1^23, 24^. It is noteworthy that exposure of DMD myotubes to agrin led to a dramatic increase in the expression of Ca_V_1.1, coinciding with the disappearance of Ca^++^ spikes. This observation suggested a potential regulatory role for agrin in modulating Ca_V_1.1 expression and subsequently influencing calcium homeostasis also in dystrophic muscle cells.

Therefore, both dystrophin and agrin, independently from each other, play a crucial role in preventing Ca^++^ spikes and regulating the expression of the Ca^++^ sensor Ca_V_1.1. It is likely that agrin, like dystrophin, stabilizes dystroglycan (DGK) on the membrane, which may either directly or indirectly contribute to maintaining Ca_V_1.1 within the complex. The addition of agrin to DMD cultures only moderately increases dystroglycan expression, suggesting the potential involvement of other structural proteins (Suppl Fig.3). Agrin, being a large molecule with heparan sulphate chains carrying a strong negative charge, exhibits limited diffusion from the NMJ. This restricted diffusion could explain why agrin is not present in sufficient quantities to stabilise DGK in the absence of dystrophin and away from a genetically corrected nucleus. For this reason, we generated a lentivector expressing mini-agrin, a small and diffusible molecule but still able to perform its function. Mini-agrin not only tabolishes Ca^++^ leakage *in vitro* but it also restores the expression of Ca_V_1.1 and sarcoglycan in vivo. The data presented here allow us to propose a tentative model that is illustrated in Fig.5. It is well-established that DHPR and RyR1 form a complex that regulates Ca^++^ fluxes from internal stores. In the absence of dystrophin, reduced expression of Ca_V_1.1 disrupts DHPR function, leading to dysregulated Ca^++^ fluxes. Re-expression of dystrophin or ectopic expression of mini-Agrin restores Ca_V_1.1 expression and abolishes Ca^++^ transients, thereby reconstituting, at least in part, the dystrophin-associated glycoprotein complex (DGC). Notably, a recent study demonstrated that increased agrin expression in skeletal muscle reverses age-associated sarcopenia in mice^25^, further supporting the potential therapeutic applications of agrin. Future studies will investigate whether co-expression of mini-Agrin and small nuclear RNAs (snRNAs) engineered to induce exon skipping in the dystrophin gene may synergistically enhance their therapeutic efficacy, in consideration of the limited intra-cellular diffusion of a snRNA in comparison with the presumed higher diffusion of a small peptide in the space between the sarcolemma and the endomysium. Experiments in large animals will conclusively address this issue.

**Fig. 5:**
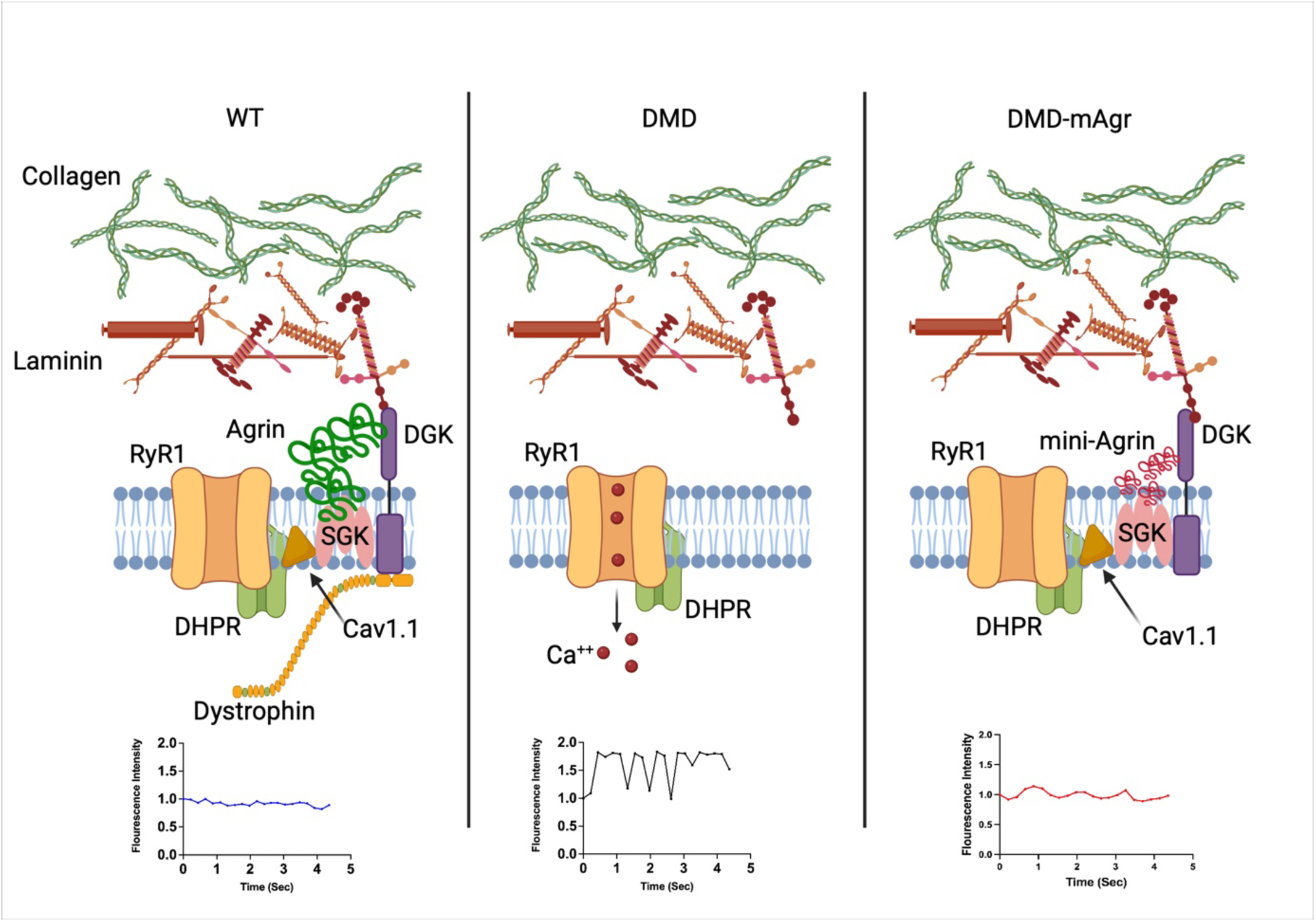
Proposed Mechanism of Mini-agrin action in Restoring Ca_V_1.1 Expression in DMD Myotubes.

## ACKOWLEDGEMENTS

This work was supported by MRC grants MR/P016006/1 and MR/S015116/1, ERC-ADG 884952 and Wellcome Trust HICF 107572.

## Materials and Methods

### Mice

The female pregnant CD1 mice and NGS-DMD mice carrying a skippable mutation of exon 51 (NOD.Cg-*Prkdc^scid^ Il2rg^tm1Wjl^* Tg (HLA-A/H2-D/B2M)1Dvs/SzJ) were obtained from Jackson Laboratory (USA) and housed in the University of Manchester animal facility. All experiments and procedures were approved by the University of Manchester Animal Welfare and Ethical Review Body (AWERB). All Experiments were conducted under the University of Manchester Animal Care guidelines and in accordance with UK Home Office Regulations (ASPA 1986) and under the project licence PDB0C0.

### Intramuscular Injection

Eight-month-old NGS-DMD mice were injected in the Tibialis Anterior (TA) with 500,000 WT human myoblasts, DMD-U7 and DMD-mAgr. The animals were culled one month after the injection.

### Cells

#### Human immortalized myogenic cells

Human immortalized WT and DMD myoblasts were generated in Vincent Mouly Lab (Institute of Myology, Paris). DMD myoblasts are characterized with Exon 51 skippable specific mutation. The DMD-U7 cells were generated in our lab by transducing DMD cells with a lentiviral vector expressing a small nuclear RNA (U7 snRNA) engineered to skip exon 51. Human immortalized muscle cells were cultured in a mixture of Dulbecco’s modified Eagle’s medium (DMEM; SIGMA, USA; D5796-500ML) supplemented with 20% RPMI media (199; Gibco, USA; 22340-020), 20% fetal bovine serum (FBS; SIGMA, USA; F9665-500ML), 50 μg/mL Gentamicin (Gent; SIGMA, USA; G1272-10ML), 25 μg/mL Fetuin (FET; Gibco, USA; PSB10052), Human Fibroblast Growth Factor (hFGF) (50 ng/ml; Gibco, USA; 17105-041), Human Epidermal Growth Factor (hEGF) (5 ng/ml; Gibco, USA; 2129284), Dexamethasone (0.2 μg/ml; SIGMA, USA; D4902-100MG), and Insulin (5 μg/ml; Gibco, USA; 51500-056 10ML).

#### Mouse primary neural cells

##### Spinal cord isolation and cell extraction

Murine embryonic spinal cords were isolated from E12.5 CD1 embryos. The isolated spinal cords were chopped into small pieces and digested in HBBS (1X; Gibco, USA; 14175-053) containing 0.24 U Dispase (2.4 U/ml; Gibco, USA; 17105-041) for 25 minutes at 37 °c in a shaking water bath. The cell suspension was then centrifuged for 10 minutes at 280xg.

##### Culture medium for MNs

A specific medium was used to maintain and culture isolated MNs. The medium was prepared as follows: the DMEM/F-12 medium (1X; Gibco, USA; 21331-020) was supplemented with 2% B-27 (1X; Gibco, USA; A35828-01), 1% L-GLUT (1X; Gibco, USA; 15140-122), 1% Penicillin/Streptomycin (P/S) (1X; Gibco, USA; 15140-122), and 10 µg/ml Mouse Neuronal Growth Factor (1X; Gibco, USA; 17105-041).

### 3D culture of Human myotubes & mouse embryonic MNs

Human immortalised myoblasts were seeded on a layer of collagen-coated plates and incubated overnight at 37 ^°c^ ^with^ 5% CO2. The following day, the growth medium was replaced with differentiation media: DMEM supplemented with 1x insulin-transferrin-selenium-X (Gibco) and 50 μg/mL Gentamicin (Gent; SIGMA, USA; G1272-10ML). On day 6 of differentiation, extracted MNs were mixed with Matrigel suspended in MN growth medium and added on top of the myotube culture.

#### Lentivector and Transduction

The lentiviral vector U7#51T2AGFP was described in Galli et al 2024 (https://pubmed.ncbi.nlm.nih.gov/38438561/).

The lentiviral vector LV-mCherry-Mini-agrin was derived from the pLV-mCherry-EF1A backbone, allowing co-expression of mCherry and mini-agrin under the control of the CMV and human Elongation Factor-1 alpha (EF-1α) promoters, respectively (Suppl. Fig. 4A). The vectors were produced by Vector Builder (https://en.vectorbuilder.com/). Human immortalized myoblasts were transduced in T80 flasks at a virus dilution of 1:500. Forty-eight hours post-transduction, the viral medium was removed, and cells were allowed to proliferate under standard culture conditions.

#### Immunofluorescence Staining

Cells were cultured and differentiated on collagen I-coated 24-well plates (Thermo Fisher Scientific, USA; 142475), washed with PBS, and fixed in 4% paraformaldehyde (PFA) at room temperature. The cells were then permeabilised with 1% BSA in 0.2% Triton in PBS at room temperature and blocked with 10% fetal bovine serum (FBS) at room temperature. Primary antibodies used were: α-MyHC MF-20 (Development Studies Hybridoma Bank); α-Dys MANDRA17 (Development Studies Hybridoma Bank); α-TUJ1 (Biolegend, MMS-435P); α-Bungarotoxin BTX (Sigma-Aldrich, T0195), α-Ca_V_1.1 (Thermo Fischer Scientific, MA3-920); α-Sarcoglycan (Atlas, A95444); α-Dystroglycan (Invitrogen, PA5-28179); α-Serca (Invitrogen, MA3-910); α-nNOS (Invitrogen, PA3-032A). Secondary antibody used were: Donkey α-Mouse (Thermo Fischer Scientific, A21203), Goat α-Rabbit (Thermo Fischer Scientific, A11037). Nuclei were stained with Hoechst stain (Thermo Fischer Scientific, H3570). Images were acquired using a (ZEISS AX10) inverted microscope and processed and analysed using (Zeiss Zen Imaging software).

Tibialis Anterior (TA) samples were embedded in OCT (Thermo Fisher Scientific, USA; LAMB/OCT) and frozen in isopentane. 10-micron sections were fixed in 4% paraformaldehyde (PFA) at room temperature. The sections were then permeabilised with 1% BSA in 0.2% Triton in PBS at room temperature and blocked with 10% fetal bovine serum (FBS) at room temperature. Primary antibodies used were: α-Ca_V_1.1 (Thermo Fischer Scientific, MA3-920); α-Dystrophin (Merckmillipore, 574777); α-mini-Agrin (in house, https://www.biorxiv.org/content/10.1101/2025.09.16.676550v1); α-Lamin A/C (Invitrogen, MA3-1000); α-Sarcoglycan (Atlas, A95444); α-Dystroglycan (Invitrogen, PA5-28179);. Secondary antibody used were: Donkey α-Mouse (Thermo Fischer Scientific, A21203), Goat α-Rabbit (Thermo Fischer Scientific, A11037). Nuclei were stained with Hoechst stain (Thermo Fischer Scientific, H3570). Images were acquired using a (ZEISS AX10) inverted microscope and processed and analysed using (Zeiss Zen Imaging software).

### Live-Cells Ca^+*+*^ imaging

Intracellular calcium levels were measured using the green-fluorescence calcium indicator Fluo-4 AM (Invitrogen, USA, F14201). Differentiated myotubes were loaded with 2 μM of Fluo-4 AM prepared in DMSO. The cells were incubated for 1 hour at 37 °c, followed by washing with the differentiation medium. Myotubes were maintained at 37 °c, and images were captured using an inverted fluorescence microscope (ZEISS AX10) at 4.2 frames per second with 488 nm laser line excitation. The analysis was carried out using ImageJ software by measuring the fluorescence intensity of the acquired images over time and normalising the values to the fluorescence intensity at the initial time point.

### Transmission electron microscope (TEM)

WT, DMD, and DMD-U7 myoblasts were plated on collagen-I-coated ACLAR embedding film overnight at 37 ^°c^ and 5% CO₂. After ten days of differentiation, myotubes were fixed with a fixative solution containing 4% PFA, 2.5% GA, and 0.1M HEPES. Samples were processed at the Bioimaging Core facility at the University of Manchester.

### Calcium Blocking Treatments

Nimodipine (NIM, Sigma, USA, N-149) was dissolved in DMSO and used at 10 μM. Nitrendipine (NIT, Sigma, USA, N-149) was dissolved in DMSO and used at 10 μM. Differentiated myotubes were treated with either NIM or NIT for 30 minutes prior to live imaging. Caffeine (Sigma, USA, C-0750) was dissolved in dH2O and used at 10 mM. All the drugs were added to fully differentiated myotubes during the Ca^+*+*^ imaging.

### Agrin treatment

Human recombinant agrin powder (50 μg; R&D Systems, USA; 6624-AG) was reconstituted with PBS and used at the required concentration for each experiment.

### qRT-PCR

RNA was extracted from differentiated myotubes using TRIzol reagent (Life Technologies, USA, 15596018) and purified according to the manufacturer’s protocol. cDNA was synthesised using a cDNA preparation kit (Thermo Fisher Scientific, USA).

For qRT-PCR reactions, the sequences of the primer pairs were as follows:

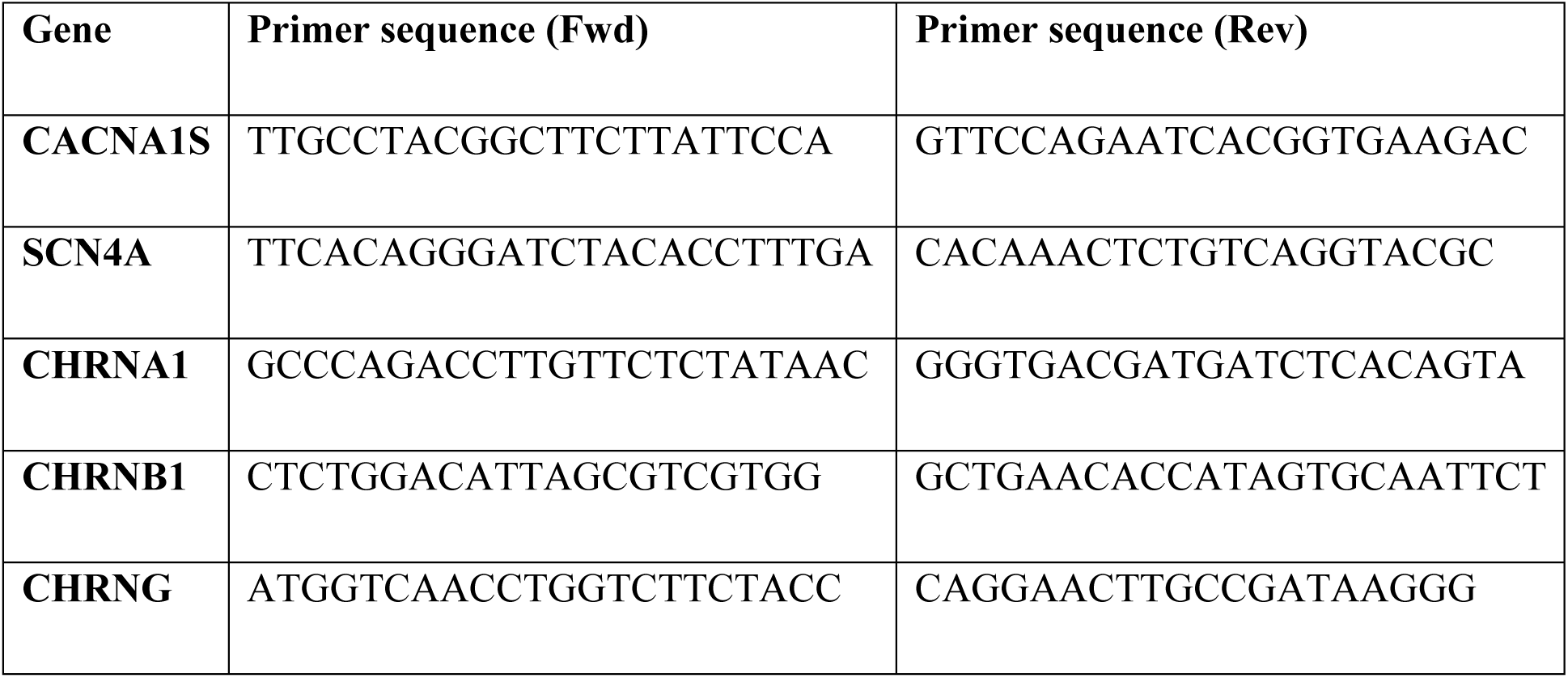
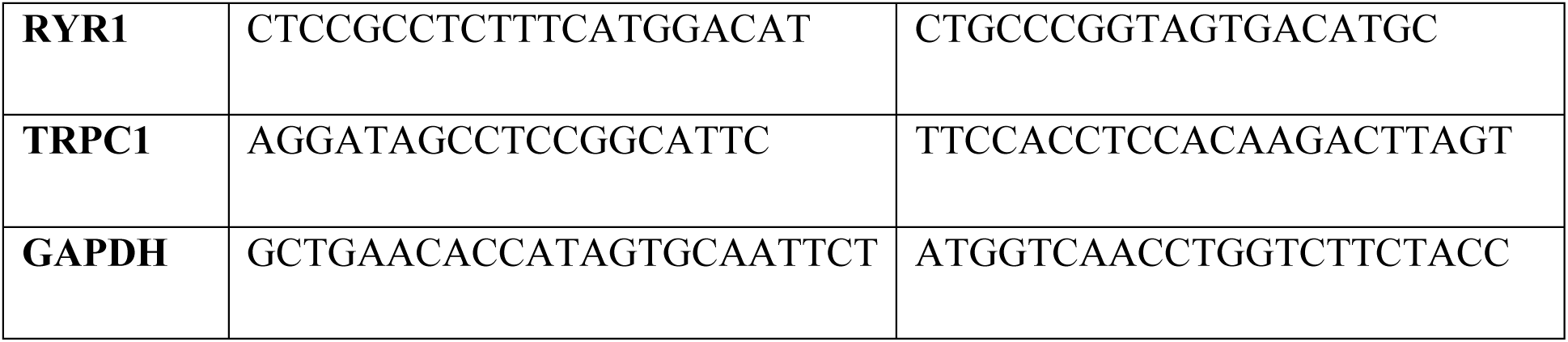

### Western Blot

Cells were lysed in RIPA buffer (10mM TRIS, 100 mM NaCl, 1mM EDTA, 1% Triton, 10% Glycerol, 0.1% SDS and 1% protease inhibitor (78446, Thermo Fisher Scientific, USA). Protein concentration was determined with the Bio-Rad Protein Assay. Proteins were separated in 10% gradient, pre-casted SDS PAGE and then transferred onto a nitrocellulose membrane using standard protocol. The blots were incubated with the following antibodies: α-Ca_V_1.1 (Thermo Fischer Scientific, MA3-920); α-Sarcoglycan (Atlas, A95444); α-Dystroglycan (Invitrogen, PA5-28179); α-Vinculin (Abcam, V9131); α-GAPDH (Abcam, Ab 125247). Secondary antibody used was: HRP conjugated mouse IgG (Dako, P0447); HRP conjugated rabbit IgG (Dako, P0448).

Proteins were visualized by an enhance chemiluminescence method (Thermo Fisher Scientific, 34580) according to the manufacturer’s instructions.

### Statistical Analysis

All in vitro experiments were repeated at least three times, each in triplicate, to ensure robust and reproducible measurements. For qRT-PCR, RNA gene expression was normalized to GAPDH, and comparisons between groups were performed using the Welch t-test in GraphPad Prism (v8.4.2) to account for unequal variances. In vivo experiments were conducted on experimental groups of three animals, representing a balance between the 3R principles and the need for reproducibility and consistency. Sample size was determined based on previous experimental data, expected effect size, and variability, using a Z-score of 1.96 for 95% confidence, a population standard deviation of 0.5, and an effect size of 0.5, resulting in a minimum of three animals per group to achieve sufficient statistical power. All data are presented as mean ± standard deviation (SD). Differences were considered statistically significant at p < 0.05.

