## Supplementary Figure for "Mini-agrin prevents calcium leakage and restores the dystrophin complex"

### Supplementary Figures

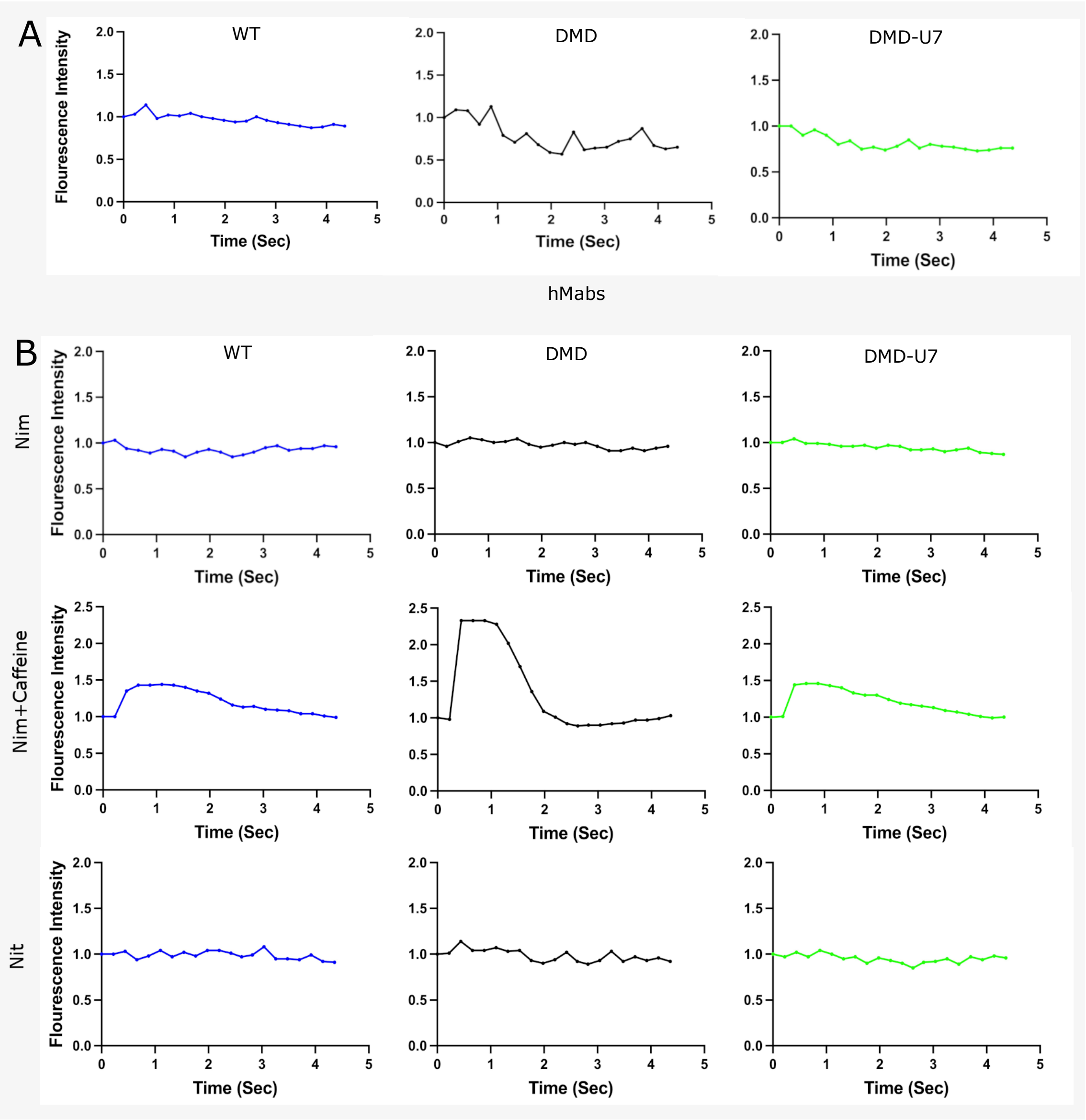


**Fig. S1: Aberrant $Ca^{2+}$ Spikes in DMD Myotubes Are Mediated by $Ca_V1.1$ Channels.**

**A.** Representative fluorescence intensity (FI) analysis over time in WT, DMD, and DMD-U7 hMABs. Values were normalized to FI at time 0 (n = 3). **B.** Representative FI analysis over time in WT, DMD, and DMD-U7 myotubes treated, from top to bottom, with the calcium channel blocker nimodipine (Nim, 10 mM), nimodipine (Nim, 10 mM) plus caffeine (10 mM), or nitrendipine (10 mM). Values were normalized to FI at time 0 (n = 3).

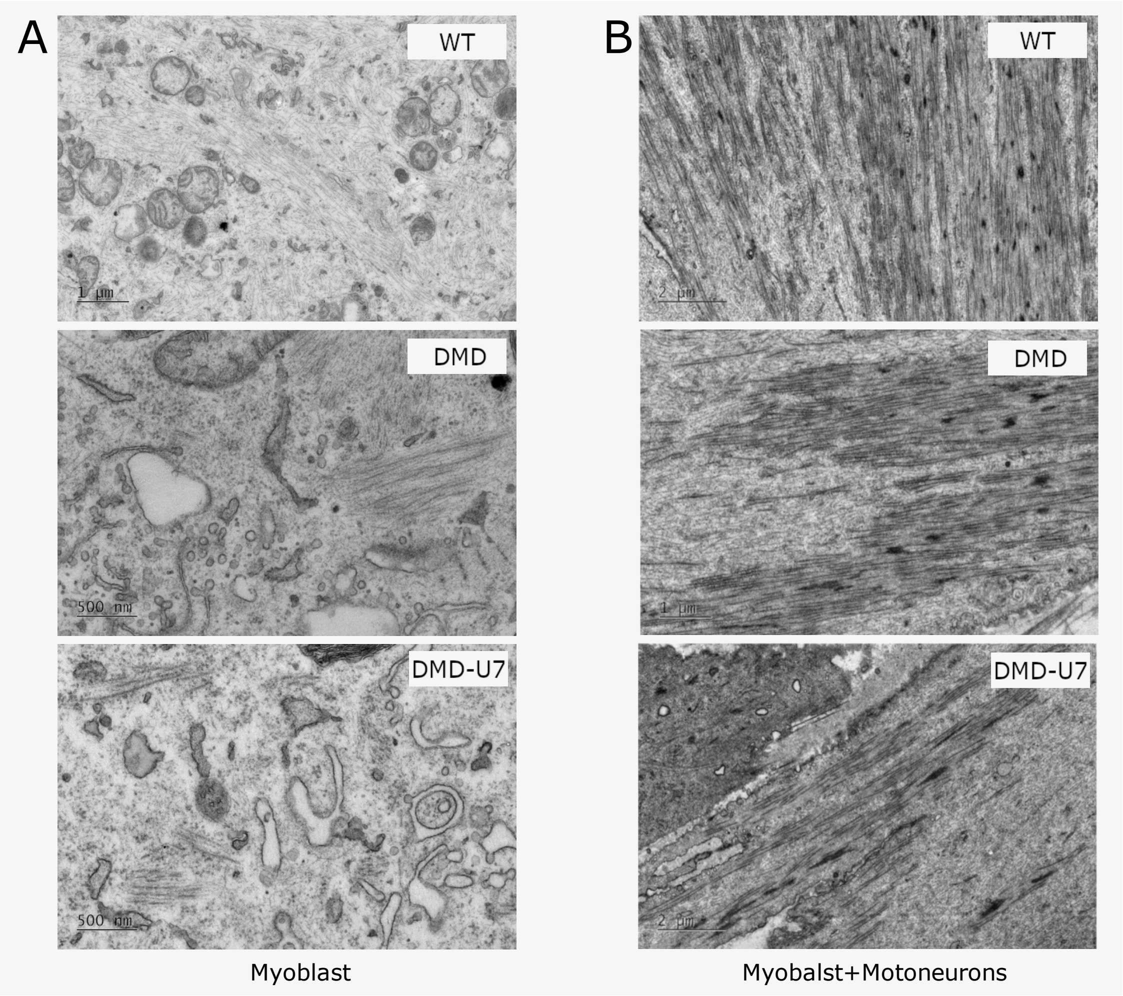


**Fig. S2: Motoneuron Co-culture Promotes Maturation and NMJ Formation in DMD Myotubes.**

**A.** Representative TEM image of early-differentiated WT, DMD, and DMD-U7 myotubes (n=3). **B.** Representative TEM image of early-differentiated WT, DMD, and DMD-U7 myotubes co-cultured with mouse motoneurons (n=3).

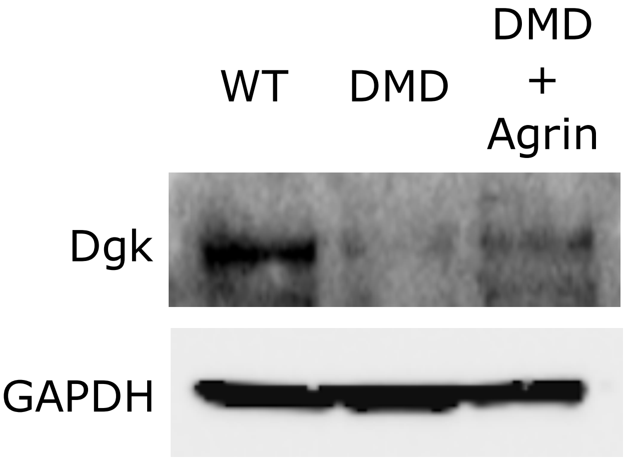


**Fig. S3: Agrin Restores Dystroglycan Expression in DMD Myotubes.**

Representative WB showing dystroglycan (Dgk) expression in WT, DMD, and DMD myotubes treated with agrin (7 µg/ml) (n=3).

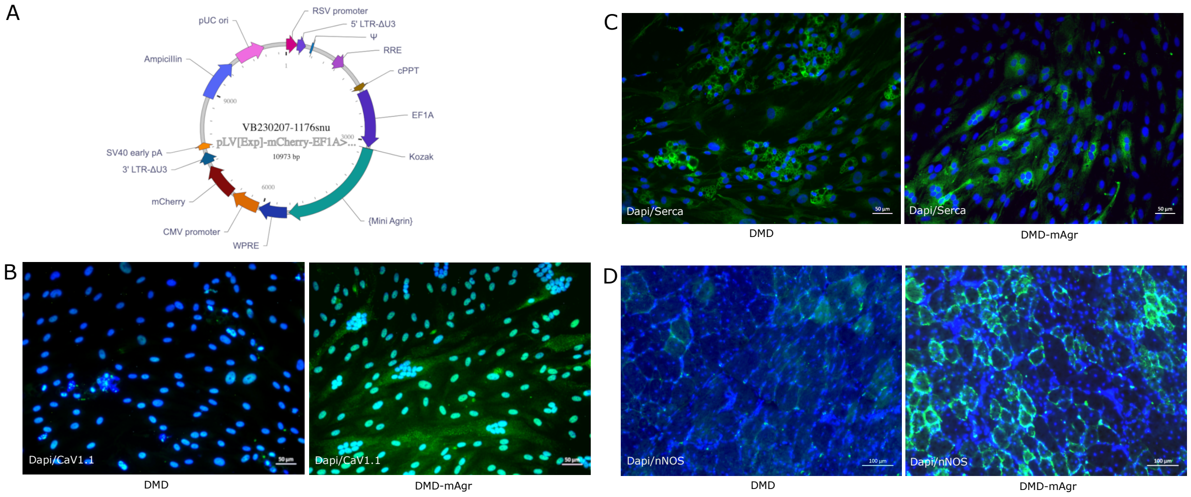


**Fig S4: Mini-agrin Expression Restores Key Muscle Proteins in DMD Myotubes and Muscle.**

**A.** Schematic map of the lentiviral vector expressing mini-agrin. **B.** Immunofluorescence (IF) images of DMD and DMD-mAgr myotubes showing $Ca_V1.1$ expression (green) (n=3). **C.** IF images of DMD and DMD-mAgr myotubes showing SERCA expression (green) (n=3). **D.** IF images of transverse sections of tibialis anterior (TA) muscles from NSG-mdx-Δ51 mice transplanted with $5 \times 10^5$ DMD-mAgr cells, showing nNOS expression (green) (n=3).
